# *Drosophila* Modeling of Insomnia-Associated Genes Reveals Diverse Underlying Sleep Phenotypes

**DOI:** 10.1101/2025.06.12.659376

**Authors:** Torrey R. Mandigo, Farah Abou Daya, Suraj Math, Athena Q. Ye, Matthew Maher, Richa Saxena, James Walker

## Abstract

Insomnia is a prevalent sleep disorder with highly heterogenous manifestations. While data-driven approaches to insomnia subtyping have revealed potential differences between proposed insomnia subtypes and their impacts on overall health, little is known about the genetic factors that underly and differentiate these potential insomnia subtypes. We utilize a human-genetics driven approach to *Drosophila* modeling to identify the range of sleep traits regulated by insomnia-associated genes. Modeling pan-neuronal loss of *Drosophila* orthologs of a set of insomnia genes reveals a broad range of sleep phenotypes. Through systematic characterization of traits related to sleep quantity, timing, and quality, we identify genetic factors that co-regulate aspects of the insomnia-associated phenotypic landscape. Out of the 75 insomnia-associated genes identified, only 1/3 had at least one *Drosophila* ortholog that regulated overall sleep quantity. In contrast, 1/3 of the insomnia-associated genes had at least one *Drosophila* ortholog that regulated either sleep timing or sleep quality, without impacting sleep quantity. Together this work, in *Drosophila*, provides support for a genetic influence on the differences between insomnia subtypes.

## Introduction

Insomnia is the most common sleep disorder [1] and can be characterized by a range of sleep complaints including issues falling asleep, maintaining sleep, waking up too early with difficulty going back to sleep, or overall sleep issues impacting daytime functions [2–5]. The heterogeneity of insomnia manifestations likely contributes to the 6% to 50% range in prevalence found across epidemiological studies, along with the varied definitions used in each study [6]. While twin studies have found the heritability of insomnia to range from 38-59% [7], recent large-scale genome-wide association studies (GWAS) have implicated an abundance of genetic loci that contribute to risk of developing insomnia [8,9]. However, the varied manifestations of insomnia suggest that variants in these loci may lead to a range of sleep phenotypes underlying the broader classification of insomnia.

There has long been debate around the potential for clinically relevant insomnia subtypes [10]. Efforts in data-driven subtyping has been attempted to better address the heterogeneity within insomnia disorder [11,12]. Studies investigating differences between insomnia subtypes have even found deviations in structural brain connectivity and attentional brain responses among suggested subtypes [13,14]. Furthermore, exploratory assessment of sleep restriction therapy, on insomnia patients, indicates subtypes with altered slow wave activity may see better improvements in sleep measures than those with normal slow wave activity [15]. Insomnia subtypes can also have varied impacts on the risk of health conditions such as pain [16], mood disorders [17], stroke [18], and cardiovascular disease [19]. While the benefits of recognizing insomnia subtypes is becoming clearer, the molecular mechanisms and genetic factors underlying each subtype remains poorly understood.

Here, we reveal a broad range of sleep phenotypes caused by the neuronal disruption of insomnia-associated genes, mirroring differences seen in some potential insomnia subtypes. Through *Drosophila* modeling, we characterize the impacts on measures of sleep quantity, timing, and quality for *Drosophila* orthologs of insomnia-associated genes identified from human GWAS. Investigation of genetic regulators of sleep quantity revealed subsets of genetic factors that coregulate activity levels during wake periods and those that impact sleep without altering wake activity. Characterization of sleep timing revealed genetic factors that distinctly impact daytime sleep or nighttime sleep with varied effects on overall sleep quantity. Lastly, characterization of sleep quality, through measures of sleep fragmentation and continuity, found genetic factors that resulted in fragmented sleep and overall sleep loss as well as factors that lead to sleep fragmentation with no overall impact on sleep quantity. In total, our characterization of the role of insomnia-associated genes in regulating varied aspects of sleep reveals co-regulators of sleep quantity, timing, and quality and independent regulators of distinct sleep traits further supporting the potential for insomnia subtypes at the genetic level.

## Methods

### Orthology

Predicted *Drosophila* orthologs were determined using the DRSC integrative ortholog prediction (DIOPT) tool [20]. Weighted DIOPT scores were collected using DIOPT version 8.0. Genes were included in the list of *Drosophila* orthologs with human associations to insomnia if they were the highest scoring ortholog for an insomnia-associated human gene or if their weighted DIOPT score was within 1 of the top scoring ortholog. Upon the release of DIOPT version 9.0 weighted DIOPT scores were re-evaluated for any unscreened genes and removed from the list of they no longer fit the previously mentioned criteria. A reciprocal *Drosophila* ortholog was defined as a *Drosophila* gene with the best human to *Drosophila* weighted score (Best Score) and the best *Drosophila* to human weighted score (Best Reverse Score). Non-reciprocal *Drosophila* orthologs were defined as a *Drosophila* gene with the best human to *Drosophila* weighted score (Best Score) but a had a closer ortholog in the human genome when assessing orthology from *Drosophila* to human.

### Drosophila Husbandry

All *Drosophila* stocks used in this study were raised on standard cornmeal fly food under a 12h/12h light cycle at 25°C and 50-75% relative humidity. Fly stocks used in this study were obtained from Bloomington *Drosophila* Stock Center (BDSC) including the pan-neuronal Gal4, *elav-Gal4* (BDSC#: 458), and control lines, *UAS-GFP RNAi* (II) (BDSC#: 41552) and *UAS-GFP RNAi* (III) (BDSC#: 41553), or from Vienna *Drosophila* Resource Center (VDRC), including control lines, GD (VDRCC#: 60000) and KK (VDRC#: 60100).

### Sleep Assays and Analysis

The GAL4-UAS system was used to drive knockdown of our genes of interest in a tissue specific manner. Flies carrying a UAS-RNAi construct were crossed to *elav-Gal4* flies and incubated at 25°C. Adult F1 male progeny were collected. Sleep-wake behavior was collected through an IR beam break method using the *Drosophila* Activity Monitoring (DAM) system (Trikinetics, Waltham MA). Individual 1-3-day post-eclosion male flies were loaded into glass tubes containing the same cornmeal fly food stocks were raised on. Flies were allowed to adjust and entrain to monitoring conditions and the monitoring incubator light sources (12h/12h light/dark cycle, 25°C, and 50-75% relative humidity) for 2 days. After entrainment, activity data was collected for 5 days. Flies that died during the 5-day data collection were excluded from our analysis. Beam-break records of activity were analyzed using ClockLab and RStudio. Periods of sleep were defined as any bout of inactivity lasting five minutes or longer as previously described [21,22]. Measure of total sleep includes any sleep during the 24-hour light cycle (ZT0-ZT24). Wake activity accounts for the average amount of beam breaks per minute awake. Day sleep includes any sleep during the light period of the light cycle (ZT0-ZT12), while night sleep includes any sleep during the dark period of the light cycle (ZT12-ZT24). Anticipatory behavior was calculated as a ratio between the total activity levels in the 3-hour period preceding a light transition compared to the total activity in the 6-hour period preceding the same light transition, as previously described [23]. Morning anticipation includes the activity levels for the transition from dark to light and evening anticipation includes activity levels for the transition from light to dark. Sleep latency represents the amount of time between the transition to the dark phase of the light cycle and the first bout of sleep. In the case where a fly was asleep during the light to dark transition, no sleep latency was calculated for that day. Bout number represents the total number of discreet periods of sleep during one 24-hour light cycle (ZT0-ZT24), while bout length represents the average length of the discreet periods of sleep during one 24-hour light cycle (ZT0-ZT24). Sleep discontinuity represents the number of one-minute wake periods, the minimal possible amount of time between sleep bouts, during the 24-hour light cycle. All measurements of sleep and activity metrics represent averages from the 5 days of monitoring. At least 20 flies of each genotype were analyzed from at least 2 independent experiments. Any genotypes with less than 20 flies represent genotypes with reduced viability. In these cases, the total amount of flies that survived the 5-day sleep-wake monitoring, from at least 4 independent experiments, were presented. All experiments were conducted on male flies to avoid complications and disruptions in activity monitoring caused by egg laying and wander larva leading to erroneous beam breaks during data collection. Beam break counts from DAM systems was analyzed using custom R scripts. RStudio scripts and methodology can be found on https://github.com/jameswalkerlab/Gill_et.al..

### Statistical Analysis

For all sleep-wake behavior metrics, statistical significance was determined by one-way analysis of variance (ANOVA) followed by Šidák’s multiple comparisons post hoc test comparing each genotype to their respective control. P-values (Pval) used for Δ Sleep, Δ Wake Activity, Δ Day Sleep, Δ Night Sleep, Δ Bout Number, or Δ Bout Length are the adjusted P-values from Šidák’s multiple comparisons post hoc tests. All bar graphs show mean ± SD, unless otherwise stated. All statistical analyses were performed with GraphPad Prism 9.

## Results

### Identification of insomnia associated genes and their Drosophila orthologs for characterization of sleep traits

We began by identifying insomnia-associated genes to model in *Drosophila* to understand the phenotypic landscape and the genetic factors regulating various sleep traits, that may be driving insomnia associations (**Fig 1**). First, we curated an initial set of 69 insomnia-associated genes from gene mapping efforts by Jansen *et al*., utilizing positional mapping, eQTL mapping, and/or chromatin interaction mapping, or from annotations by Lane *et al*. identifying a given gene as the only gene within an associated locus [8,24]. Additionally, we included a smaller set of 6 genes gathered from single gene association peaks for other sleep traits, including napping, short sleep, excessive daytime sleepiness, and sleepiness, of which 4 of the 6 were found to also share associations with insomnia [8,9]. From our subset of 75 insomnia-associated genes, we next identified orthologs in the *Drosophila* genome using DRSC integrative ortholog prediction tools (DIOPT). Altogether, we collated 93 *Drosophila* orthologs of the 75 human genes and performed RNAi-mediated pan-neuronal knockdown of each, driven by *elav-Gal4*. In total, we screened a 282 RNAi lines, including lines from the P-element RNAi (GD) and phiC31 RNAi (KK) libraries from the Vienna *Drosophila* Stock Center (VDRC) and the TRiP collection from the *Drosophila* RNAi Screening Center. Adult flies with knockdown of each insomnia-associated ortholog were evaluated for various aspects of sleep quantity, timing, and quality to provide a characterization of the impact of each gene on the regulation of sleep.

**Figure 1.**
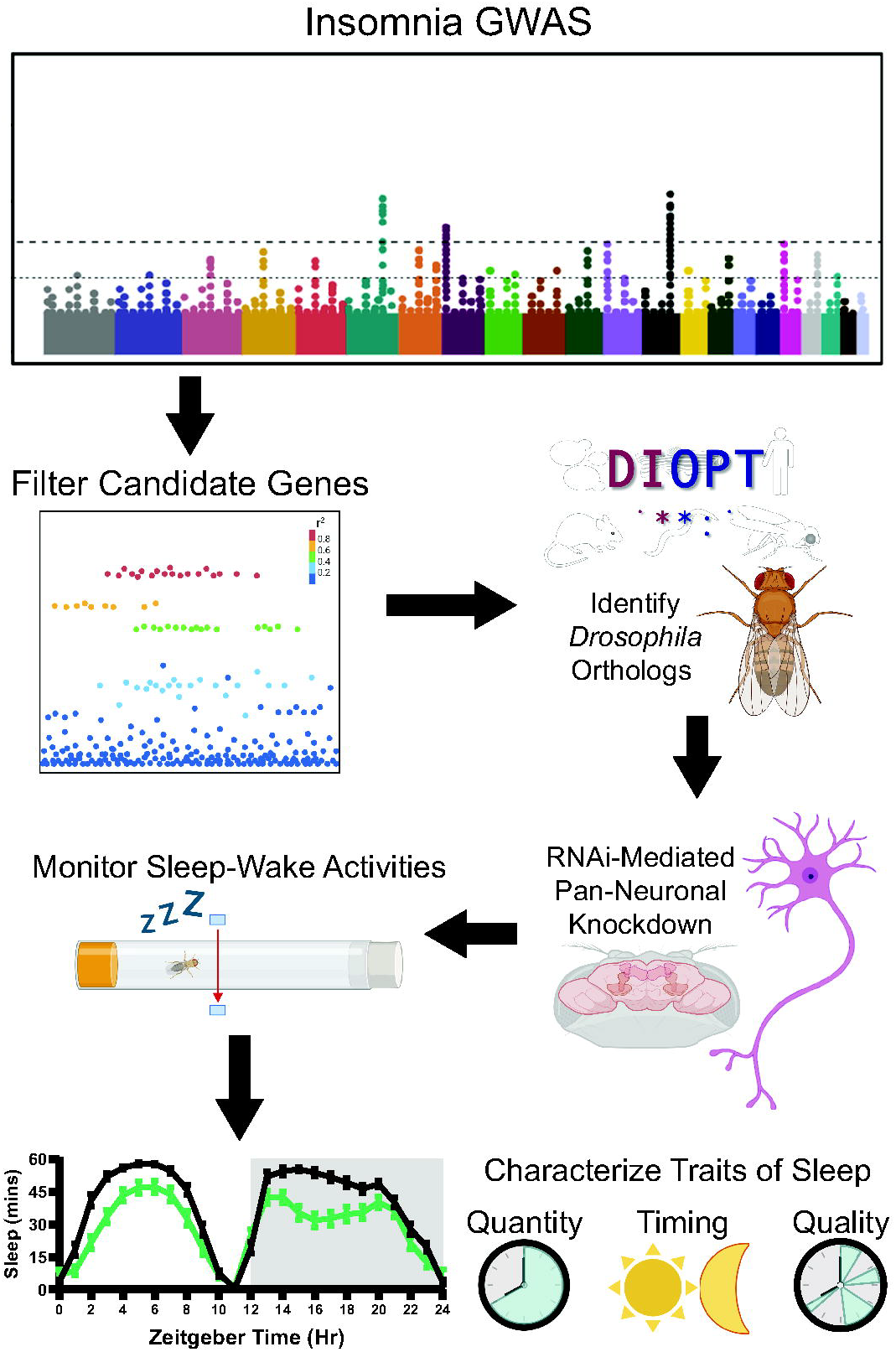
Experimental pipeline from identifying insomnia-associated genes to characterization in *Drosophila*. Graphical schematic of the workflow pipeline for identifying and prioritizing human insomnia-associated genes from GWAS. Insomnia associated genes were filtered and *Drosophila* orthologs were identified. RNAi-mediated pan-neuronal knockdown was performed and impacts on sleep traits related to quantity, timing, and quality were characterized.

### *Drosophila* modeling of insomnia-associated genes reveals varied relation between sleep quantity and wake activity

We assessed which *Drosophila* orthologs of genes in insomnia loci, from human GWAS, functioned in regulating sleep quantity. Of the 93 orthologs, 55% (51 genes) showed some evidence of regulating sleep quantity with at least one RNAi line impacting total minutes slept. Notably of the 282 RNAi lines tested, only two resulted in a significant increase in sleep quantity, while 77 resulted in a significant decrease. 23 orthologs demonstrated robust evidence of regulating sleep quantity with two or more RNAi lines resulting in a significant decrease in total minutes slept. Furthermore, no *Drosophila* genes had multiple RNAi lines that significantly increased sleep. Since sleep and activity are intricately linked, we examined how the rate at which insomnia associated genes co-regulate sleep quantity and activity levels when awake. Of 282 RNAi lines, 53 (19%) resulted in altered sleep quantity with no impact on wake activity, while 52 lines decreased sleep and one increased sleep. Conversely, 38 RNAi lines (13%) resulted in changes in wake activity with no impact on sleep quantity, 37 lines decreased wake activity and one increased activity. 26 RNAi lines (9%) led to a change in both sleep quantity and wake activity, 22 lines decreased both sleep and activity, three lines decreased sleep and increased activity, and one increased both sleep and activity (**Fig 2A,B,C**).

**Figure 2.**
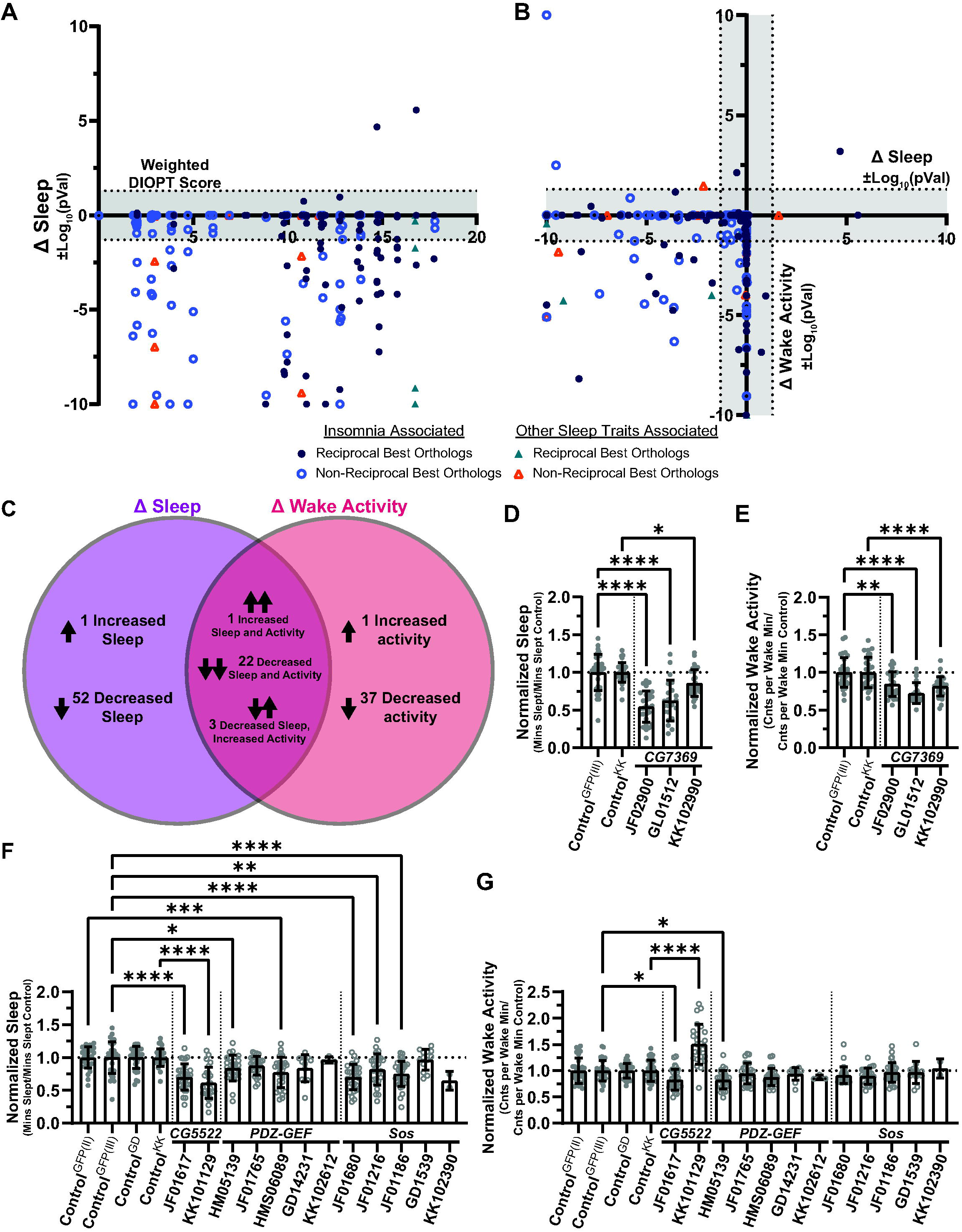
Insomnia-associated genes can have varied impacts on the sleep quantity and activity levels. (**A**) Distribution of the impacts on sleep quantity, of *Drosophila* lines targeting orthologs of insomnia-associated genes, by their weighted DIOPT score. (**B**) Comparison of impacts on sleep quantity and wake activity for each *Drosophila* line tested. (**A,B**) Individual dots represent the log10(p-value) of the change in the represented parameter compared to a respective control (averaged over 5 days) with negative indicating a decrease and positive indicating an increase. Shaded areas designate statistically non-significant (p>0.05) log10(p-value). Triangles indicate points with values greater than the specified scale ranges. (**C**) Venn Diagram summarizing data from *Drosophila* lines impacts on sleep quantity and wake activity. Changes are only reported for lines with statistically significant impacts on sleep or wake activity. (**D-G**) Quantification of (**D,F**) total sleep and (**E,G**) wake activity for a select set of *Drosophila* orthologs of RasGefs. Each dot represents the mean for an individual fly over 5 days. Each data point is normalized to the mean of the respective control for each RNAi. Error bars represent standard deviation. Adjusted p-values from Šídák’s multiple comparisons test are indicated with asterisks (*p<0.05, **p<0.01, ***p<0.001, ****p<0.0001).

Next, we investigate the consistency of the relationship between the regulation sleep quantity and activity levels within insomnia-associated genes that encode proteins with similar functions. When assessing positive regulators of total minutes slept, identified by our *Drosophila* modeling, we found that orthologs of multiple RasGEFs regulated the amount of sleep flies experienced in a 24-hour period. Pan-neuronal knockdown of *CG7369*, an ortholog of *RASGEF1B* (DIOPT_v9.0_ weighted score: 16.75; Reciprocal), resulted in a significant decrease in total minutes slept compared to controls for all three RNAi lines tested (*elav*>*CG7369*^*JF02900*^, *elav>CG7369*^*GL01512*^, and *elav*>*CG7369*^*KK102990*^) (**Fig 2D**). These decreases in sleep were accompanied by reduced activity during wake periods for all three RNAi lines (**Fig 2E**). Two orthologs of another RasGEF, *RASGRP1*, similarly regulated sleep quantity. Pan-neuronal knockdown of *CG5522* (DIOPT_v9.0_ weighted score: 1.81, Non-Reciprocal), with *elav*>*CG5522*^*JF01617*^ and *elav*>*CG5522*^*KK10112*^, and knockdown of Sos (DIOPT_v9.0_ weighted score: 1.81, Non-Reciprocal), with *elav*>*Sos*^*JF01680*^, *elav*>*Sos*^*JF01216*^, and *elav*>*Sos*^*JF01186*^, all resulted in significant decreases in total sleep (**Fig 2F**). However, while knockdown of *Sos* did not impact activity levels with any RNAi lines tested, knockdown of *CG5522* resulted in highly variable impacts on wake activity. *elav*>*CG5522*^*JF01617*^ resulted in decreased activity and *elav*>*CG5522*^*KK101129*^ resulted in increased activity while awake (**Fig 2G**), suggesting the impact of *CG5522* knockdown on wake activity may be sensitive to background genetics. *Sos* also scored as an ortholog of *RASGRF2* (DIOPT_v9.0_ weighted score: 2.84; Non-Reciprocal). Similar to *Sos*, another ortholog of *RASGRF2, PDZ-GEF* (DIOPT_v9.0_ weighted score: 1.94; Non-Reciprocal), also regulated sleep quantity. Pan-neuronal knockdown of *PDZ-GEF* with *elav*>*PDZ-GEF*^*HM05139*^ and *elav*>*PDZ-GEF*^*HMS06089*^ both led to a decrease in total minutes slept (**Fig 2F**), yet only knockdown using *elav*>*PDZ-GEF*^*HM05139*^ resulted in significantly decreased wake activity (**Fig 2G**). Together these data demonstrate that even orthologs with highly similar functions such as RasGEFs can vary in their co-regulation of sleep quantity and activity levels.

### *Drosophila* modeling of insomnia-associated genes reveals regulators of sleep timing independent of changes in sleep quantity

Next, we aimed to assess which *Drosophila* orthologs of insomnia-associated genes regulate sleep timing. To do this we characterized each ortholog’s role in regulating daytime sleep or nighttime sleep. Of 93 orthologs tested, 35% (32 genes) had a single RNAi line and 16% (15 genes) had two or more RNAi lines that resulted in significant changes in daytime sleep. Alternatively, 34% (31 genes) had a single RNAi line and 18% (16 genes) had two or more RNAi lines that resulted in significant changes in nighttime sleep (**Fig 2A**). This suggests orthologs of insomnia-associated genes are equally likely to regulate daytime and/or nighttime sleep in *Drosophila*. To understand the relationship between the regulation of sleep quantity and sleep timing, we assessed the frequency at which changes in daytime or nighttime sleep coincided with changes in overall sleep quantity. Of the 121 RNAi lines with significant changes in day sleep, night sleep, and/or total sleep, 18% (22 RNAi lines) had changes in all three metrics. 24% (29 RNAi lines) had changes in daytime sleep and total sleep. 23% (28 RNAi lines) affected nighttime sleep and total sleep. 20% (24 RNAi lines) had changes in daytime sleep that did not result in significantly altered total sleep. While 14% (17 RNAi lines) had changes in nighttime sleep that did not result in significant changes in sleep quantity. Lastly, there was a single RNAi line that resulted in significantly altered daytime and nighttime sleep, without impacting total sleep quantity. Together these data suggest that although alterations in sleep timing more frequently coincide with changes in overall sleep levels, some genes and pathways may regulate day and/or night sleep without impacting sleep quantity (**Fig 3A,B**).

**Figure 3.**
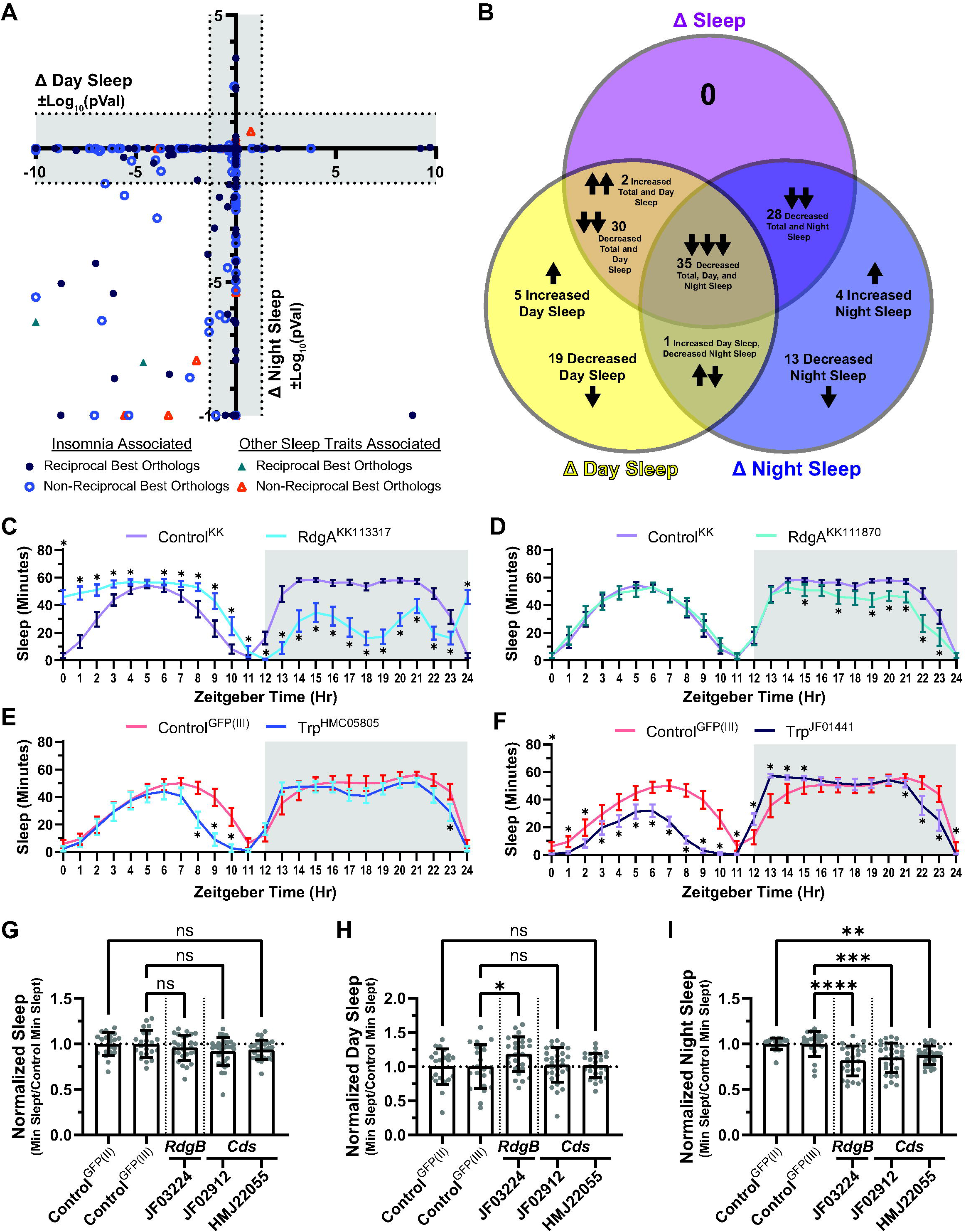
Insomnia-associated genes can regulate aspects of sleep timing independent of impacts on sleep quantity. (**A**) Comparison of impacts on day sleep and night sleep for each *Drosophila* line tested. Individual dots represent the log10(p-value) of the change in the represented parameter compared to a respective control (averaged over 5 days) with negative indicating a decrease and positive indicating an increase. Shaded areas designate statistically non-significant (p>0.05) log10(p-value). Triangles indicate points with values greater than the specified scale ranges. (**B**) Venn Diagram summarizing data from *Drosophila* lines impacts on total sleep, day sleep, night sleep. Changes are only reported for lines with statistically significant impacts on at least one the three parameters. (**C-F**) Quantification of sleep in hourly bins for pan-neuronal knockdown of RdgA or Trp, using (**C**) RdgA^KK113317^, (**D**) RdgA^KK111870^, (**E**)Trp^HMC05805^, or (**F**) Trp^JF01441^ RNAi lines. Error bars indicate 95% confidence interval for each hourly bin. Statistical significance was determined by multiple unpaired t-tests with an FDR of 1% and bins that are statistically significant from the respective control are indicated by asterisks. (**G-I**) Quantification of (**G**) total sleep, (**H**) day sleep, and (**I**) night sleep for flies with pan-neuronal knockdown of the indicated gene using the indicated RNAi line. Each dot represents the mean for an individual fly over 5 days. Each data point is normalized to the mean of the respective control for each RNAi. Error bars represent standard deviation. Adjusted p-values from Šídák’s multiple comparisons test are indicated with asterisks (ns p>0.05, *p<0.05, **p<0.01, ***p<0.001, ****p<0.0001).

To further investigate insomnia-associated genes that function in regulating sleep timing independent of sleep quantity, we focused on the single RNAi line that impacted both day and night sleep without significantly altering total minutes slept. This particular RNAi, *rdgA*^*KK113317*^, targets a *Drosophila* ortholog of DGKI (DIOPT weighted score: 11.9; Non-Reciprocal). Pan-neuronal knockdown of *rdgA*, with *elav*>*rdgA*^*KK113317*^, significantly increased daytime sleep and decreased nighttime sleep (**Fig 3C**). Furthermore, pan-neuronal expression of a second RNAi *rdgA*^*KK111870*^, resulted in a significant yet modest decrease in nighttime sleep without impacting daytime or total sleep (**Fig 3D**). Previous work found that light-sensitive Ca^2+^ channels encoded by *Trp* were constitutively active in *rdgA* mutants [25]. *Trp* was also identified as the highest scoring ortholog of *TRPC7* (DIOPT weighted score: 4.81; Non-Reciprocal), which was included in our set of insomnia-associated gene identified from human GWAS. Pan-neuronal knockdown of *Trp*, with *elav*>*Trp*^*HMC05805*^, resulted in a modest yet significant decrease in daytime sleep with no impact on overall sleep (**Fig 3E**). However, knockdown of *Trp*, with *elav*>*Trp*^*JF01441*^, resulted in a significant decrease in daytime sleep leading to a decrease in overall sleep (**Fig 3F**). Together these data suggest that some regulators of sleep timing that may function through pathways and mechanisms that are also able to regulate overall sleep quantity.

rdgA encodes a diacylglycerol kinase that functions in phospholipase C signaling and phosphoinositide recycling. RdgB similarly functions in phosphoinositide recycling and encodes a phosphatidylinositol transfer protein that has been shown to regulate night sleep [26]. To further test the role of this pathway in regulating sleep timing, independent of sleep quantity, we assessed knockdown of *rdgB* and *Cds*, which encoded CDP-diacylglycerol synthase and functions in phosphoinositide recycling, for their role in regulating total, daytime, and nighttime sleep. Pan-neuronal knockdown of *rdgB* or *Cds* did not significantly affect total minutes slept (**Fig 3G**). However, knockdown with *elav*>*rdgB*^*JF03224*^ resulted in a significant increase in day sleep and significant decrease in night sleep (**Fig 3H,I**), similar to *rdgA*^*KK113317*^-mediated knockdown, while knockdown of *Cds*, with *elav*>*Cds*^*JF02912*^ and *elav*>*Cds*^*HMJ22055*^, both significantly decreased nighttime sleep without altered daytime sleep (**Fig 3H,I**), similar to *rdgA*^*KK111870*^-mediated knockdown. Together these data further demonstrate a role for phosphoinositide recycling in regulating sleep timing independent of sleep quantity.

### *Drosophila* modeling of insomnia-associated genes uncovers candidates that solely regulate sleep quality, independent of sleep quantity

Given our human genetics-driven approach to *Drosophila* modeling uncovered regulators of sleep quantity and sleep timing, we next evaluated which *Drosophila* orthologs of insomnia-associated genes regulate sleep quality. To assess sleep quality, we focused on sleep structure characterized by changes in sleep bout number and average sleep bout length. Out of the 93 *Drosophila* orthologs tested, 39% (36 genes) had a single RNAi line and 26% (24 genes) had two or more RNAi lines that significantly impacted sleep bout number. Of the 282 RNAi lines tested, none resulted in a significant decrease in sleep bout number, while 91 RNAi lines lead to a significant increase in sleep bouts. Furthermore, we found 40% (37 genes) had a single RNAi line and 43% (40 genes) had two or more RNAi lines that significantly impacted sleep bout length. Of the RNAi lines tested, only two resulted in a significant increase in sleep bout length, while 136 resulted in a significant decrease in bout length. Since fragmented sleep is a common manifestation of insomnia, characterized by an increase in sleep bouts that are shorter in length, we next examined how many of the regulators of sleep quality demonstrated a sleep fragmentation phenotype. We found that 38% (35 genes) had a single RNAi and 23% (21 genes) had at least two RNAi lines that resulted in sleep fragmentation (**Fig 4A**). Next, we wanted to know the rate at which fragmented sleep coincides with changes in sleep quantity. Of the 159 RNAi lines with significantly fragmented sleep and/or altered total sleep, 22% (35 RNAi lines) had both significantly fragmented sleep and decreased sleep quantity, 28% (45 RNAi lines) had significantly fragmented sleep but no impact on sleep quantity, 8% (12 RNAi lines) had significantly decreased sleep with no sleep fragmentation, and one line had increased sleep with no change to sleep structure (**Fig 4B**). Together these data suggest that sleep fragmentation can occur at similar rates with and without overall impacts on sleep quantity.

**Figure 4.**
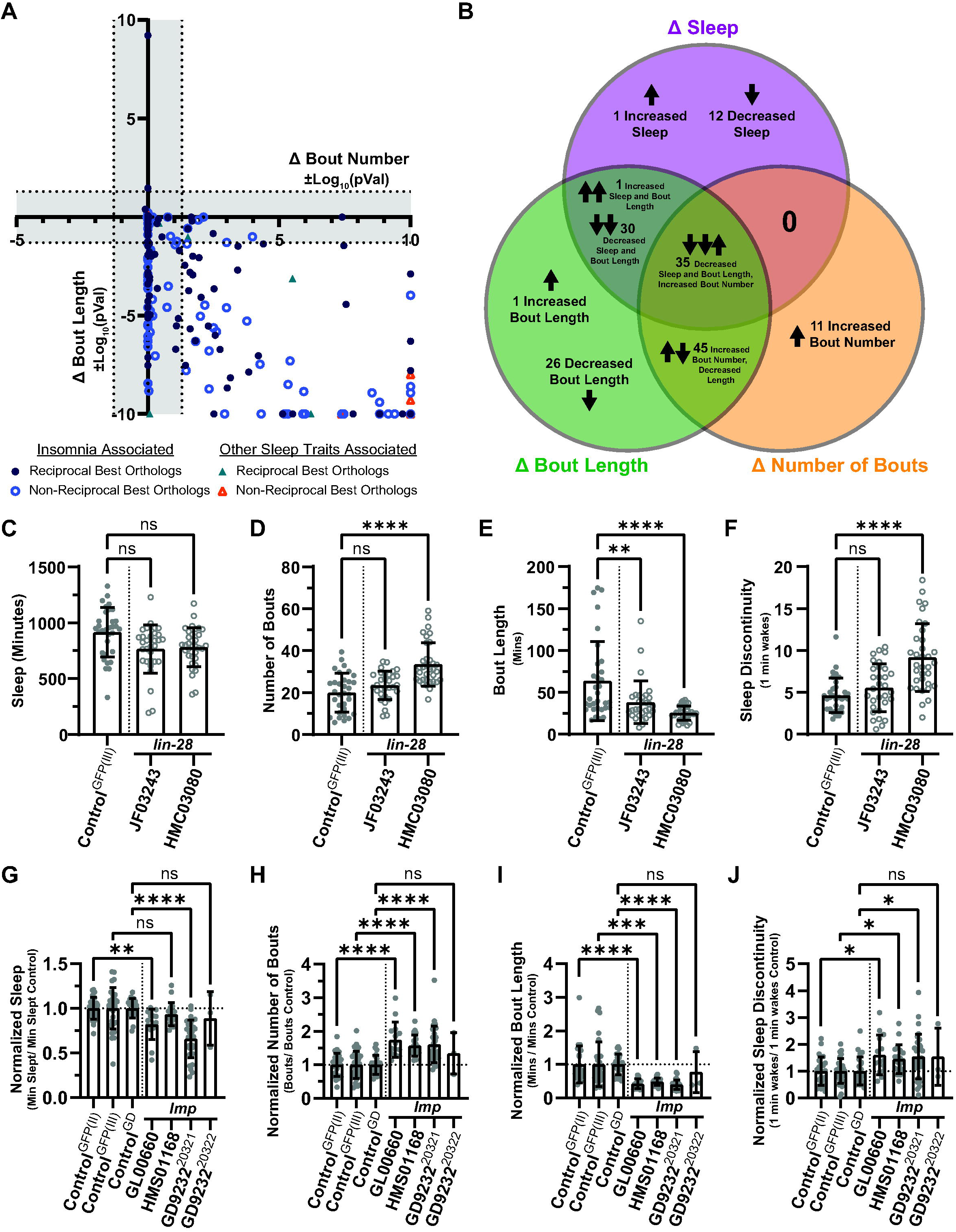
Insomnia-associated genes can regulate sleep quality and structure without impacting sleep quantity. (**A**) Comparison of impacts on sleep bout number and bout length for each *Drosophila* line tested. Individual dots represent the log10(p-value) of the change in the represented parameter compared to a respective control (averaged over 5 days) with negative indicating a decrease and positive indicating an increase. Shaded areas designate statistically non-significant (p>0.05) log10(p-value). Triangles indicate points with values greater than the specified scale ranges. (**B**) Venn Diagram summarizing data from *Drosophila* lines impacts on total sleep, sleep bout number, and bout length. Changes are only reported for lines with statistically significant impacts on at least one the three parameters. (**C-J**) Quantification of (**C,G**) total sleep, (**D,H**) number of sleep bouts, (**E,I**) sleep bout length, and (**F,J**) Sleep Discontinuity for flies with pan-neuronal knockdown of the indicated gene using the indicated RNAi line. Each dot represents the mean for an individual fly over 5 days. Each data point is normalized to the mean of the respective control for each RNAi. Error bars represent standard deviation. Adjusted p-values from Šídák’s multiple comparisons test are indicated with asterisks (ns p>0.05, *p<0.05, **p<0.01, ***p<0.001, ****p<0.0001).

To further investigate insomnia-associated genes that function in regulating sleep quality independent of sleep quantity, we focused on a high scoring, non-reciprocal ortholog, *lin-28. Lin-28* is orthologous to two paralogous human genes, *Lin28A* (DIOPT weighted score: 15.75, reciprocal) and the insomnia-associated *Lin28B* (DIOPT weighted score: 14.79, Non-Reciprocal). Pan-neuronal knockdown of *lin-28* with *elav*>*lin-28*^*HMC03080*^ resulted in significantly more sleep bouts that were overall shorter in length, without significantly altering total minutes slept Knockdown using *elav*>*lin-28*^*JF03243*^, significantly decreased the length of sleep bouts but did not impact total minutes slept or the overall number of sleep bouts (**Fig 4C,D,E**). Knockdown of lin-28 (by *elav*>*lin-28*^*HMC03080*^) also led to an increase in sleep discontinuity, measured by the number of one-minute wakes (**Fig 4F**). lin-28 was previously found to interact with Imp [27], IGF-II mRNA-binding protein, to regulate adult stem cell proliferation in the fly intestine. Therefore, we tested whether *Imp* similarly impacts sleep structure. While knockdown using two of the four RNAi lines targeting *Imp, elav>Imp*^*GL00660*^ and *elav>Imp*^*GD9232-20321*^, resulted in a significant increase in sleep fragmentation and sleep discontinuity while also causing a decrease in total minutes slept, one RNAi line, *elav>Imp*^*HMS01168*^, resulted in significantly increased sleep fragmentation and sleep discontinuity, without impacting overall sleep quantity (**Fig 4G,H,I,J**). Together these data highlight a regulator of sleep quality in *lin-28* that may partially function through a mechanism that regulates that also regulates sleep quantity.

## Discussion

In this study, we have characterized the disruption of *Drosophila* orthologs of insomnia-associated genes identified in human GWAS to uncover a range of sleep phenotypes underlying the broad insomnia classification. Focusing on a curated set of genes from insomnia human GWAS, we performed pan-neuronal knockdowns of *Drosophila* orthologs and screened for sleep phenotypes. Characterization of sleep quantity found robust evidence for a significant change in sleep amount, in at least one *Drosophila* ortholog, for only 1/3 of the insomnia-associated human genes. Expanding our characterization of sleep phenotypes to metrics of sleep timing, we found that 13 of the 75 human genes had at least one ortholog that regulated aspects of sleep timing without effecting sleep quantity. Similarly, we found 12 of the 75 human genes had at least one ortholog that regulated characteristics of sleep quality without impacting sleep quantity. We further highlight RasGEFs as a group of sleep quantity regulators with varied impacts on activity levels during wake periods, phospholipase C signaling and phosphoinositide recycling as a pathway regulating sleep timing independent of sleep quantity, and *lin-28* as a regulator of sleep quality. Together these findings highlight genetic mechanisms that regulate the range of the phenotypic landscape found in insomnia. We also identify genetic factors that regulate discrete characteristics of sleep that may contribute to differences seen in potential insomnia subtypes.

### Genetics of sleep and contribution of *Drosophila* modeling to our mechanistic understanding

In the last decade, major advances have been made in uncovering genetic variants and risk loci associated with various sleep traits and disorders through human GWAS [8,9,24,28–39]. However, in many cases these association studies reveal variants and loci for which the underlying causal gene and mechanisms remain poorly understood. In these cases, further studies in model organisms can provide valuable insight. In particular, the genetics tools available and rapid lifespan of *Drosophila melanogaster* provides a prime model for studies interrogating causal genes and mechanisms underlying these associations. Since the identification of a sleep-like state in *Drosophila* [21,22], it has been well established that mechanisms regulating sleep are highly conservated from flies to humans [40]. Genetic variation in *Drosophila* has even been harnessed, through the *Drosophila* Genetics Reference Panel (DGRP), to perform sleep related GWAS in flies [41–48]. Furthermore, SNPs associated with measures of sleep and circadian rhythm in *Drosophila* are near genes with human orthologs that similarly share sleep trait associations from human GWAS. In our set of insomnia-associated genes alone, 22 of the 93 *Drosophila* orthologs were also near SNPs with sleep trait associations in *Drosophila* GWAS [45– 48]. While many of these orthologs functioned in regulating similar characteristics of sleep between the identified SNPs and our RNAi-mediated knockdown approach, some genes with no effects on any measured sleep traits, in this study, were found to be associated with changes in circadian period or day-to-day variability. For example, no ortholog of the insomnia associated gene *SAMD5* significantly impacted any of the sleep metrics we tested. However, *SKIP*, an ortholog of *SAMD5*, was found in a circadian-focused *Drosophila* GWAS to regulate the consistency of daily activity patterns, indicating that some insomnia associations, like that of *SAMD5*, may be driven by underlying circadian disruptions not investigated in our study [46]. Conversely, many genes with insomnia associations in humans, have yet to be associated with sleep phenotypes in flies, either through *Drosophila* GWAS studies or directed investigations. Nonetheless, further work combining the genetic associations from large scale human association studies and the experimental tools in model organisms, such as *Drosophila*, will be essential for understanding the causal mechanisms and support the identification of underlying sleep phenotypes.

### *Drosophila* modeling can facilitate the delineation of potential insomnia subtypes

Insomnia is the most common sleep disorder occurring in 10-20% of adults [49]. However, insomnia is a broad classification that can include difficulty initiating sleep, maintaining sleep, experiencing non-restorative sleep, or waking up too early impacting overall daily function [50,51]. As the number of studies focusing on insomnia increases, the potential for insomnia subtype classifications has gained appreciation [14,52–54]. In our *Drosophila* modeling, only 1/3 of our human genes had at least one *Drosophila* ortholog that regulated total sleep amount. While studies focused on short sleeping insomnia-like phenotypes in flies have provided invaluable insight regarding sleep regulatory processes [55–59], it is becoming clearer that insomnia-associations can be driven by phenotypes other than short sleep. For example, a variant-to-gene mapping study of insomnia loci uncovered a conserved role of GPI-anchor biosynthesis in regulating sleep. However, disruption of the insomnia associated gene *PIG-Q* in flies and zebrafish both led to significantly increased sleep [60]. Some work has already been conducted regarding daytime sleepiness loci to identify potential biological subtypes, with risk alleles clustering predominately with either sleep propensity or sleep fragmentation phenotypes. Moreover, the sleep fragmentation subtype of these daytime sleepiness alleles was associated with increased insomnia symptoms [38]. Furthering our understanding of potential insomnia subtypes could also provide a clearer insight into risks associated with discrete sleep phenotypes underlying the broad insomnia classification. Recent work from our group, found that long or excessive sleep caused by disruption of insomnia-associated *Drosophila* orthologs more often resulted in altered heart rate. Similarly, traits related to insomnia with a focus on increased sleep, correlate with adverse cardiovascular outcomes in humans [19]. However, the number of studies focusing on mechanisms driving insomnia, in the absence of altered sleep quantity, remain lacking. Importantly, our *Drosophila* modeling found that insomnia-associated human genes were equally likely to have at least one ortholog regulating sleep timing and/or quality independent of sleep quantity, as they were to have at least one ortholog regulating sleep quantity. While independent regulators of sleep timing or quality remain understudied, one study using *Drosophila* to interrogate a single insomnia associated locus, found that the cardiac-specific disruption of some genes in this locus caused increased sleep fragmentation without impacting sleep quantity. Importantly, this work indicates that some of these regulators of sleep timing or sleep quality can even function from outside the brain [61]. Nonetheless, while experiments in model organisms unquestionably enhance our understanding of causal mechanisms and underlying sleep phenotypes driving insomnia, some advances can also be made in humans. As the use of large-scale human studies helps to drive sleep research, increasing the specificity of questions asked and data gathered through questionnaires is one avenue to that could help further our ability to delineate insomnia subtypes toward more personalized therapeutic approaches.

### Conclusions and Limitations of the Study

In summary, our study has modeled the disruption of *Drosophila* orthologs of genes with insomnia associations from human GWAS to reveal a range of underlying sleep phenotypes. However, some limitations are important to recognize when interpreting our findings. Firstly, our modeling only knocked down the fly orthologs in neurons; however, many other tissues and cell types are known to influence sleep. Additionally, we only performed RNAi-mediated knockdown, yet some variants could result in gain of function or increased expression. Our work also only focused on characteristics of sleep quantity, timing, and quality under 12h:12h Light:Dark conditions, while it is possible some of these genes may function in regulating circadian rhythms, homeostatic sleep, or more complex sleep phenotypes which were not examined in our characterizations. Given the large number of genes in this study, we restricted our experiments to male flies to avoid complications surrounding egg lying during the sleep monitoring period. However, some genes may function in regulating sleep in a sex-specific manner or play a greater role in regulating sleep in females compared to males. Lastly, we tested the highest scoring orthologs for each of the insomnia associated genes in our set but some of these orthologs had very low orthology and therefore our finds may not be as translational to human sleep as some of the higher scoring orthologs. Together these limitations are important to address in future studies, particularly for the insomnia-associated genes that did not impact any of the sleep traits measured in this work.

## Acknowledgements

We would like to thank members of our labs for critical comments on the manuscript

We acknowledge funding from the following sources:

T.M.: NIH T32HL007901.

R.S., J.A.W., G.C.M.: NIH-NHLBI 1R01 HL146751-01A1

We would like to acknowledge the *Drosophila* Research & Screening Center for the DIOPT ortholog search tool. Stocks obtained from the Bloomington Drosophila Stock Center, and Vienna Drosophila Resource Center were used in this study. We would like to thank Stephanie Bouley for her help in preparing the manuscript and proof-reading.

## Author Contributions

T.R.M., R.S. and J.A.W conceived the idea and designed the experiments.

R.S. and J.A.W. provided overall project leadership and funding support.

F.A., M.M. and R.S. performed human genetics analysis and validation.

T.R.M, S.M. and A.Y. performed the behavioral assays and associated *Drosophila* genetics.

T.R.M. prepared the manuscript with input from R.S. and J.A.W.

